# Functional Characterization of Enhancer Evolution in the Primate Lineage

**DOI:** 10.1101/283168

**Authors:** Jason C. Klein, Aidan Keith, Vikram Agarwal, Timothy Durham, Jay Shendure

## Abstract

**Background:** Enhancers play an important role in morphological evolution and speciation by controlling the spatiotemporal expression of genes. Due to technological limitations, previous efforts to understand the evolution of enhancers in primates have typically studied many enhancers at low resolution, or single enhancers at high resolution. Although comparative genomic studies reveal large-scale turnover of enhancers, a specific understanding of the molecular steps by which mammalian or primate enhancers evolve remains elusive.

**Results:** We identified candidate hominoid-specific liver enhancers from H3K27ac ChIP-seq data. After locating orthologs in 11 primates spanning ~40 million years, we synthesized all orthologs as well as computational reconstructions of 9 ancestral sequences for 348 “active tiles” of 233 putative enhancers. We concurrently tested all sequences (20 per tile) for regulatory activity with STARR-seq in HepG2 cells, with the goal of characterizing the evolutionary-functional trajectories of each enhancer. We observe groups of enhancer tiles with coherent trajectories, most of which can be explained by one or two mutational events per tile. We quantify the correlation between the number of mutations along a branch and the magnitude of change in functional activity. Finally, we identify 57 mutations that correlate with functional changes; these are enriched for cytosine deamination events within CpGs, compared to background events.

**Conclusions:** We characterized the evolutionary-functional trajectories of hundreds of liver enhancers throughout the primate phylogeny. We observe subsets of regulatory sequences that appear to have gained or lost activity at various positions in the primate phylogeny. We use these data to quantify the relationship between sequence and functional divergence, and to identify CpG deamination as a potentially important force in driving changes in enhancer activity during primate evolution.

## Background

Despite seemingly large phenotypic differences between species across the primate lineage, protein-coding sequences remain highly conserved. Britten and Davidson as well as King and Wilson proposed that changes in gene regulation account for a greater proportion of phenotypic evolution in higher organisms than changes in protein sequence [1,2]. A few years later, Banerji and Moreau observed that the SV40 DNA element could increase expression of a gene independent of its relative position or orientation to the transcriptional start site [3,4]. This finding led to the characterization of a new class of regulatory elements, enhancers.

Several aspects of enhancers make them ideal substrates for evolution. Enhancers control the location and level of gene expression in a modular fashion [5]. While a coding mutation will disrupt function throughout an organism, a mutation in an enhancer may only affect the expression of a gene at a particular time and in a particular location. This modularity of regulatory elements may facilitate the development of novel phenotypes, *e.g*. by decreasing pleiotropy [6]. Enhancers also commonly exist in groups of redundant elements, referred to as shadow enhancers, which provide phenotypic robustness [7–9]. Therefore, mutations within enhancers generally exhibit lower penetrance than mutations in coding sequences, facilitating the accumulation of variation.

Researchers have studied the role of enhancers in evolution through two main methods: high-resolution, systematic analysis of single enhancers, or low-resolution, genome-wide analysis of many enhancers. Examples of the former include fruitful investigations of how specific enhancers underlie phenotypic changes, *e.g*. cis-regulatory changes of the *yellow* locus affecting Drosophila pigmentation [10,11], recurrent deletions of a *Pitx1* enhancer resulting in the loss of pelvic armor in stickleback [12], and recurrent SNPs in the intron of *MCM6*, resulting in lactase persistence in humans [13,14].

Low-resolution, genome-wide approaches for discovering candidate enhancers via biochemical marks, when applied to multiple species, have identified large-scale turnover of enhancers between human and mouse embryonic stem cells [15], human and mouse preadipocytes and adipocytes [16], Drosophila species [17], mammalian limb bud [18] and vertebrate and mammalian liver [19,20].

High-resolution studies can provide clear insights into the evolution of individual enhancers, but the findings may not be broadly generalizable. Low-resolution studies have the advantage of characterizing thousands of enhancers at a time, but fail to pinpoint functional variation. Massively parallel reporter assays (MPRAs) may offer an opportunity to bridge the insights offered by low- and high-resolution studies. MPRAs have enabled high-resolution functional dissection of enhancers by testing the effects of naturally occurring and synthetic variation on regulatory activity [21–26], but have not yet been extensively applied to study enhancer evolution.

Here we set out to concurrently study the evolutionary-functional trajectories of hundreds of enhancers with MPRAs. We identified potential hominoid-specific liver enhancers based on genome-wide ChIP-seq and then functionally tested all of these in parallel. After identifying “active tiles” of these candidate enhancers, we tested eleven primate orthologs and nine predicted ancestral reconstructions of each active tile for their relative activity. Normalizing to the activity of the reconstructed sequences of the common ancestor of hominoids and Old World monkeys, we identify several subsets of active tiles that appear to have gained or lost activity along specific branches of the primate lineage; only some of these patterns are consistent with ChIP-seq-based expectations. We also use these data to examine how mutational burden impacts enhancer activity across the phylogeny, quantifying the correlation between sequence divergence and functional divergence. Finally, we examine the set of mutations that appear to drive functional changes, and find enrichment for cytosine deamination within CpGs.

### Results

#### Identification of candidate hominoid-specific enhancers

From a published ChIP-seq study in mammals [19], we identified 10,611 H3K27ac peaks (associated with active promoters and enhancers) that were present in humans and absent from macaque to tasmanian devil, and that were not within 1 kilobase (Kbp) of a H3K4me3 peak (associated with active promoters). We considered this set of peaks as potential hominoid-specific enhancers (active within the clade from gibbon to human). We narrowed this to a subset of 1,015 candidate enhancers overlapping ChromHMM strong-enhancer annotations in human HepG2 cells [27] that also had orthologous sequences in the genomes of species from human to marmoset. On average, the intersection between the hominoid-specific H3K27ac peak and HepG2 ChromHMM call was 1,138bp (**Supplemental Figure 1A**). In order to identify active subregions of each candidate enhancer, we designed 194nt sequences tiling across the length of each, overlapping by 93–100bp (**Figure 1A**).

**Figure 1.**
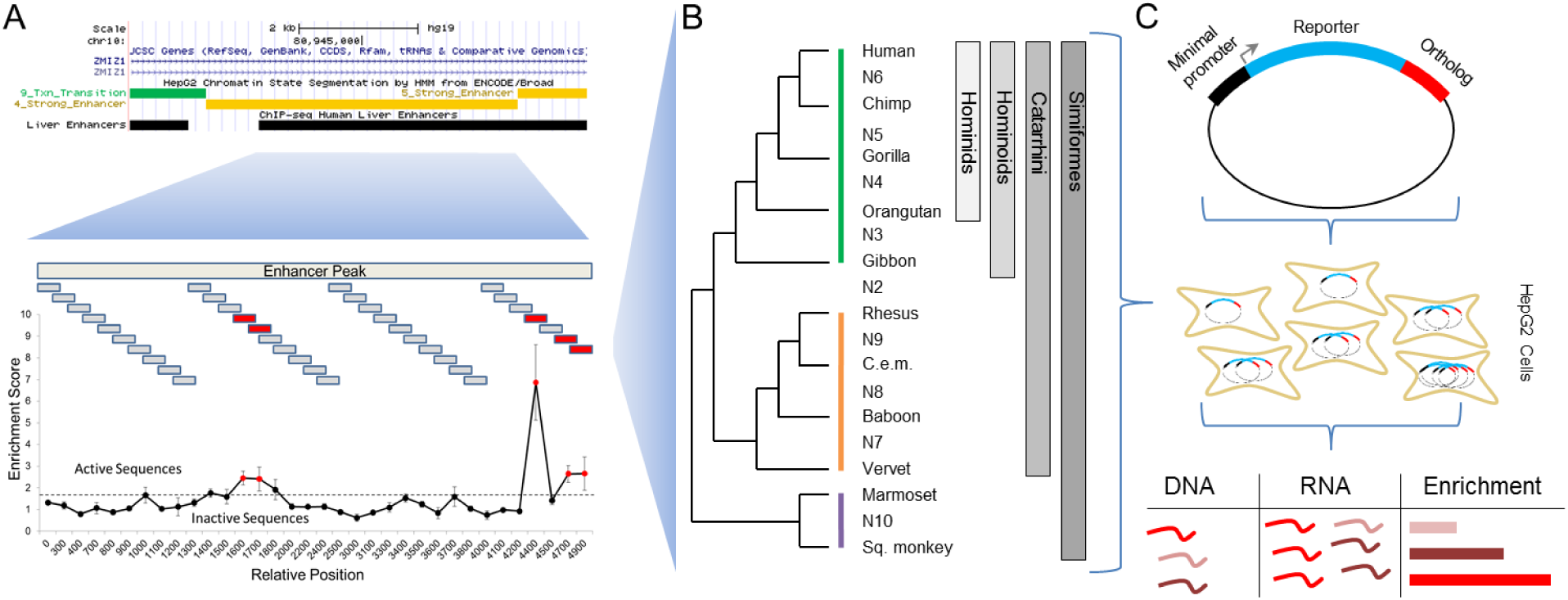
Schematic of experimental design. A) We identified potential hominoid-specific enhancers by intersecting hominoid-specific ChIP-seq predicted enhancers from primary human liver with ChromHMM-predicted strong enhancers in HepG2 cells (screenshot from http://2enome.ucsc.edu) [29]. We then tiled across each candidate enhancer using 194nt sequences and identified 697 tiles that were active in the STARR-seq reporter assay in HepG2 cells. B) We located orthologous sequences in 11 primates and computationally reconstructed 9 ancestral sequences for 348 of the active tiles. C) We then cloned all 20 present-day or ancestral orthologs per tile and performed STARR-seq again in HepG2 cells. After collecting DNA and RNA from cells, we calculated enrichment scores as the ratio of RNA to DNA for each ortholog. Each shade of red represents a different ortholog tested.

We synthesized and tested all 10,544 tiles for enhancer activity in a massively parallel reporter assay. Specifically, we used the STARR-seq vector [28], in which candidate enhancers are cloned into the 3’ UTR of an episomal reporter gene, in human HepG2 cells in triplicate. After extracting, amplifying and sequencing DNA and RNA corresponding to the enhancer region from transfected cells, we calculated an enrichment score for each tile as the ratio of RNA to DNA (rho for pairs of replicates between 0.581 and 0.676) (**Supplemental Table 1)**. We defined “active tiles” as elements with a mean enrichment score at least two standard deviations above the mean enrichment score of 125 scrambled control sequences. While most of the 1,015 candidate enhancers contained no active tiles, we identified 697 active tiles (out of 10,544, or 6.6%), occurring within 34% of the candidate enhancers (**Supplemental Figure 1B**). While we chose a strict cutoff for active tiles to increase specificity, we do note a significant shift towards more positive enrichment scores for our tiles as compared to scrambled control sequences (mean z-score 0.504 v 0.000, p<1e-5, t-test for two independent means) (**Supplemental Figure 1C**).

#### Computationally predicting the activity of ancestral and orthologous sequences

A goal of this study was to characterize how mutational burden and spectrum relate to the functional divergence of enhancer activity in primates. We used eleven high-quality primate genomes (human, chimpanzee, gorilla, orangutan, gibbon, rhesus, crab-eating macaque, baboon, vervet, marmoset and squirrel monkey) to locate similarly-sized orthologs of each of our 697 active human tiles. We were able to find orthologs in all eleven species for 348 of the 697 active human tiles. Since these species are separated by only ~40 million years, they retain high nucleotide identity. We sought to take advantage of this to ask whether we could computationally pinpoint the sequence changes that underlie apparent functional differences between orthologous sequences within primates. Of note, we had not yet measured the functional activity of orthologs of active tiles. Rather, we were assuming that previously observed patterns of gain/loss in H3K27 acetylation were reflective of whether particular tiles were active or inactive in each primate.

We first examined turnover of motifs of transcription factors (TFs) known to be associated with enhancer activity in HepG2: FosL2 and JunD [30]. We focused on comparing the human ortholog to the marmoset ortholog, the furthest outgroup with ChIP-seq data. We identified a modest enrichment of the AP-1 consensus motif, the motif for JunD and FosL2 binding, in the human ortholog compared to marmoset (p=0.012, Fisher’s exact). However, AP-1 site turnover could only explain 5% of the gain-of-function events predicted by H3K27ac ChIP-seq.

As a different approach, we built a computational model for predicting enhancer activity in HepG2 cells, and then sought to apply that model to the active tiles and their orthologs. Specifically, we trained a gapped k-mer support vector machine (gkm-SVM), a sequence-based classifier based on the abundance of gapped k-mers in positive and control training data, on an independent massively parallel reporter assay experiment in HepG2 cells [30,31]. We evaluated the model by predicting the enrichment scores from our tiling experiment on human orthologs, which the model had not seen during training. Although the original data was based on an entirely different MPRA assay (‘lentiMPRA’), the scores for each tile predicted from the gkm-SVM model correlated reasonably well with our enrichment scores obtained through STARR-seq in HepG2 cells (Spearman rho=0.453, p-value <1e-10) (**Figure 2A, Supplemental Table 2**). We then used the model to predict regulatory activity for the rhesus, vervet and marmoset orthologs, all of which did not have H3K27ac peaks. We expected to find lower predicted activity for these three orthologs compared to human. However, the predicted activity for the human vs. rhesus, vervet or marmoset orthologs were not significantly different (p=0.10, two-sample t-test), although it did trend in the right direction for all three comparisons (**Figure 2B**).

**Figure 2.**
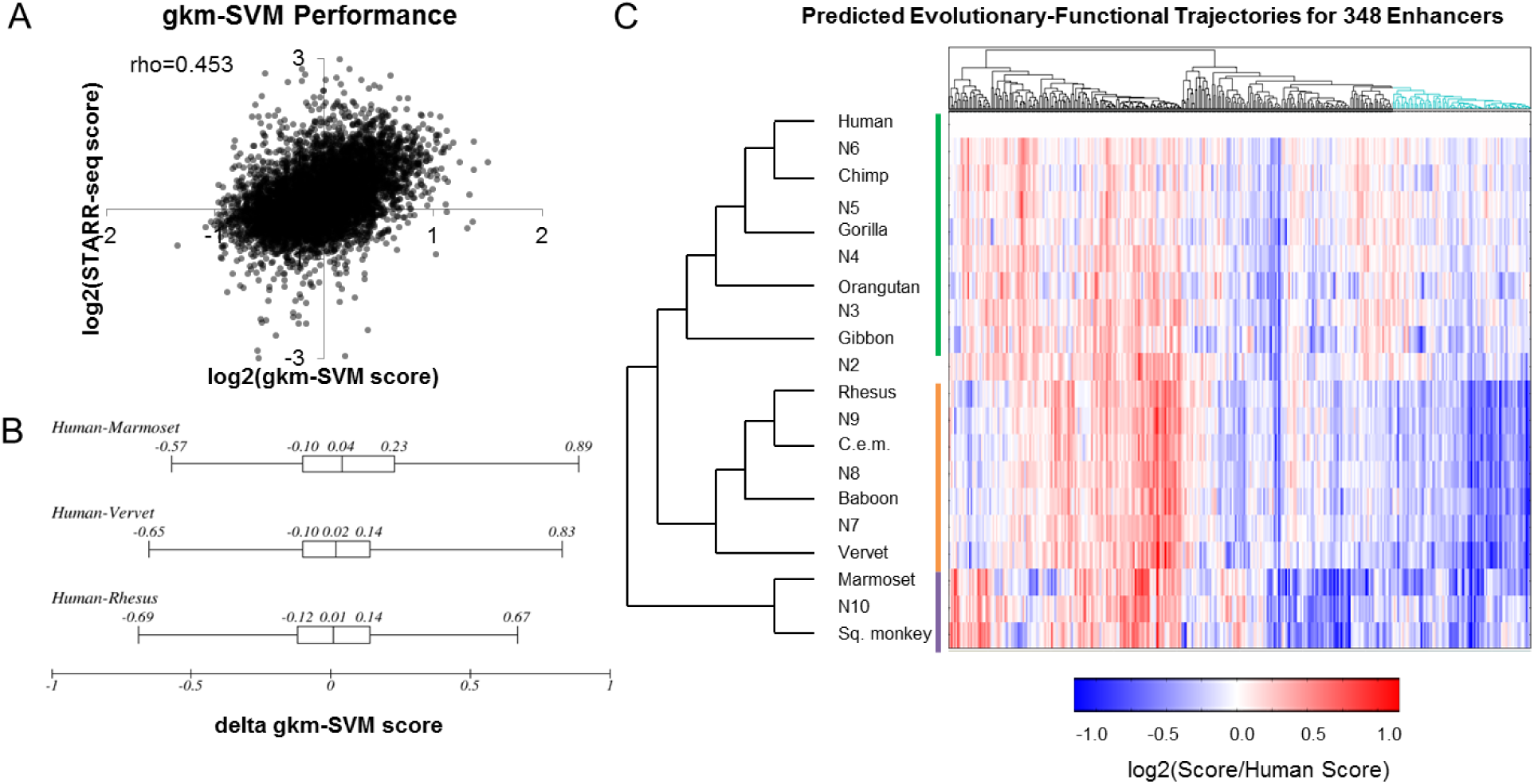
Performance of Computational Predictions. A) We trained the gapped-kmer support vector machine classifier (gkm-SVM) on an independent reporter assay experiment conducted in HepG2 cells. We then predicted the functional activity of all of our human sequence tiles and found a modest correlation with our functional data. B) The distributions of differences (delta gkm-SVM score) in predicted score between the human vs. marmoset, vervet, or rhesus ortholog for all active human tiles. C) Predicted scores for all orthologs of the 348 human-active enhancer tiles, normalized to the human ortholog. Clades are denoted by colored lines (green: hominoid, orange: Old World monkeys, purple: New World monkeys). Cyan clade of dendrogram denotes a group of 82 enhancer tiles that follows expectations for hominoid-specific enhancers as predicted by ChIP-seq comparative genomics.

With the goal of increasing our power to detect mutations underlying gains or losses in enhancer activity, we reconstructed nine ancestral sequences of the 11 primate orthologs using FastML, a maximum-likelihood heuristic (**Figure 1B**) [32]. We then applied the gkm-SVM model to predict regulatory activities for the 20 orthologs (11 from present-day primate genomes, 9 reconstructed ancestral sequences) of the human-active tiles. To characterize evolutionary trajectories, we performed hierarchical clustering on the vectors of predicted activity for each tile, and identified a group of 82 enhancer tiles that show decreased predicted activity in rhesus, vervet and marmoset compared to human, following the pattern predicted by H3K27ac ChIP-seq (**Figure 2C**). However, this was a clear minority of all tiles evaluated with this computational model (82/348 or 24%).

#### Functional characterization of ancestral and orthologous sequences

We were surprised that only a quarter of our computational predictions were concordant with ChIP-seq predictions. This could be due to limitations either in interpreting patterns in H3K27ac gain/loss, or of the computational models that we are applying to predict the relative activities of orthologs, or both. To investigate this further, we synthesized and functionally tested all 20 versions of each of the 348 active tiles with the STARR-seq vector in HepG2 cells. With the goal of improving accuracy and reproducibility, we added degenerate barcodes adjacent to each sequence of interest while cloning the library, so that we could distinguish multiple independent measurements for each element. Furthermore, we performed three biological replicates, which correlated well (independent transfections; Spearman rho between 0.773 and 0.959) (**Supplemental Figure 2A-C**). We took the average enrichment score of all barcodes over all three replicates and filtered out any element with less than six independent measurements (**Supplemental Table 3**). On average, this set had 31 independent measurements per element (**Supplemental Figure 2D**).

The resulting dataset included enrichment scores for 5,426 of the 6,960 sequences tested (78.0%), corresponding to 344 of the 348 human-active enhancer tiles (98.9%). As expected, the average pairwise correlation between species was higher within clades (hominoid, Old World monkeys and New World monkeys) than between clades (**Figure 3A**). For our initial analyses, we normalized the enrichment scores for all non-human orthologs to the enrichment score of the human ortholog, given that these tiles were first identified on the basis of the human ortholog exhibiting activity.

**Figure 3.**
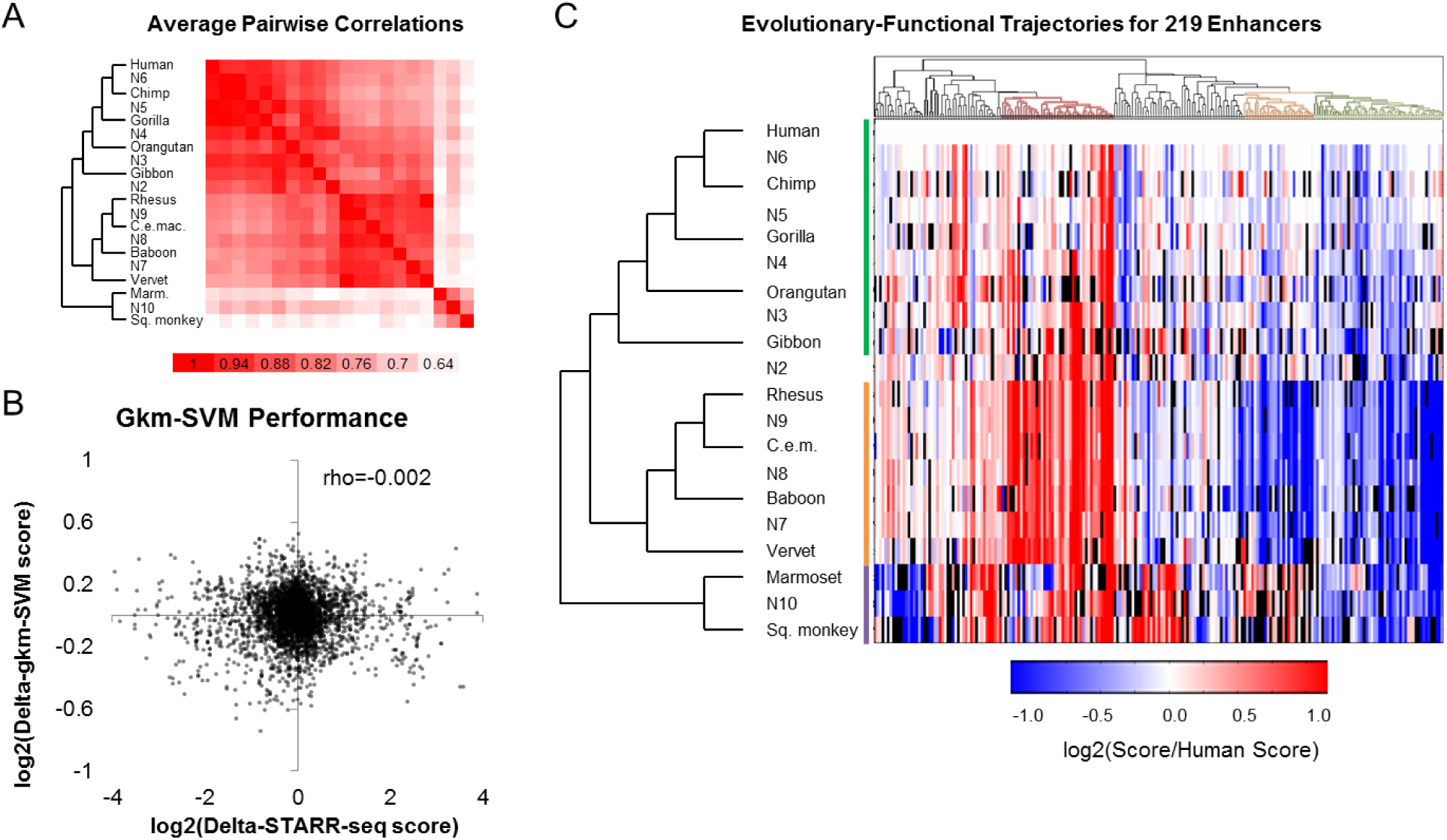
Functional scores for orthologs and ancestral sequences. A) The average pairwise Spearman correlation of functional scores between any two orthologs across all enhancer tiles tested. B) Correlation between the STARR-seq enrichment scores, normalized to the enrichment score of its human ortholog (log2[non-human score/human score]), and gkm-SVM predicted scores, similarly normalized to the predicted score of its human ortholog (log2[non-human prediction/human prediction]). C) Functional scores normalized to human for all orthologs of the 219 enhancer tiles. Black bars represent missing data. Clades are denoted by colored lines (green: hominoid, orange: Old World monkeys, purple: New World monkeys). Groups are color-coded in the dendrogram; red: relatively higher activity in in Old World monkeys, orange: relatively lower in Old World monkeys, green: relatively higher activity in either humans or hominoids.

We identified 219 enhancer tiles for which we successfully assayed activity for the human ortholog and at least 14 other orthologs. For each of these orthologs, we asked how well the experimental measurements correlated with the gkm-SVM predictions from **Figure 2C**. Specifically, we asked whether the gkm-SVM model predicted functional differences between closely related orthologs by comparing our scores of model predictions vs. functional data (all scores normalized to the human ortholog). There was no correlation between the predicted vs. experimental normalized scores (**Figure 3B**; Spearman rho= -0.002, p-value=0.892) (**Supplemental Table 4**). Therefore, while the kmer-based model performed well at characterizing relative activities of diverse elements (**Figure 2A**), it did not predict the relative activities of closely related sequences as measured here (**Figure 3B**).

We next performed hierarchical clustering on the vectors of experimentally measured activity for each tile *(i.e*. where each vector consists of the set of activities experimentally measured for all orthologs or ancestral reconstructions of a human-active tile, normalized against the activity of the human ortholog; **Figure 3C**). We identified a group of 50 enhancer tiles with relatively higher activity in either humans or hominoids (50/219 or 23%) (**Figure 3C**, green group on dendrogram), a group of 32 enhancer tiles with relatively lower activity in the Old World monkey lineage (14.6%) (**Figure 3C**, orange group on dendrogram), and a group of 43 enhancer tiles with relatively higher activity in the Old World monkey lineage (19.6%) (**Figure 3C**, red group on dendrogram). As a negative control, when we permuted species’ ids for each tile *(i.e*. shuffling raw scores represented within each column of **Figure 3C**, renormalizing, and hierarchical clustering), we no longer observe coherent clustering of activity patterns by clades (**Supplemental Figure 3**).

The first group, *i.e*. the subset of 50 enhancer tiles (23% of 219 tested) with greater activity in humans or hominoids relative to other primates, corresponds to the pattern predicted by the ChIP-seq data, and a similar proportion to the 82 tiles (24% of 348 tested) whose computationally predicted activity was concordant with the pattern predicted by the ChIP-seq data (**Fig. 2C**). However, only 14 enhancer tiles overlapped between these groups, which is not more than expected by chance (p=0.48, Fisher’s exact test). This was consistent with the lack of correlation between the experimental and predicted relative scores shown in **Figure 3B**.

For several reasons, we chose to move forward with the experimentally measured activities of primate enhancer tile orthologs. First, we believe that experimental measurements are preferable to computational predictions when available. A condition for this preference is that the experimental measurements are reproducible, which in this case they are (**Supplemental Figure 2A-C**). Second, the computational model used here is predicting the likelihood of a sequence belonging to an active vs. inactive group, while the experimental data measure the relative activity of each sequence. Experimental data is therefore better suited for quantifying differences in activity, which is the attribute that we would like to correlate with sequence divergence. Third, the differences in experimentally measured activity between orthologs relative to human were generally much greater in magnitude than the computational predictions *(e.g*. compare **Figure 2C** vs. **Figure 3C**, which use the same color scale), and furthermore in patterns that were consistent with the phylogeny relating those sequences to one another (**Figure 3C** vs. **Supplemental Figure 5**, which is the same data permuted).

### Evolutionary-functional trajectories for hundreds of enhancer tiles across the primate phylogeny

We had originally normalized enhancer tile activity to the human ortholog with the assumption that most enhancer tiles would be hominoid-specific based on patterns in H3K27ac ChIP-seq data. While our largest group did agree with the ChIP-seq data, it only represented 23% of the tested tiles. Given that the groups that we did observe were relatively coherent in relation to the lineage tree (**Figure 3C**), we turned to asking whether we could quantify the enhancer activity of various orthologs relative to their common ancestor.

For this, we normalized the enhancer tile activity scores for all orthologs to the most recent common ancestor (MRCA) of Catarrhines (N2; common ancestor of hominoids and Old World monkeys). We then performed hierarchical clustering on the 206 enhancer tiles with scores for N2 and at least 14 additional orthologs (**Supplemental Table 5**). The resulting heatmap is shown in **Figure 4A**. We observed several subsets of enhancer tiles that exhibited gains or losses in activity as measured by STARR-seq, relative to the experimentally measured activity of the reconstructed sequence of their common ancestor (**Supplemental Table 6**). Many of these subsets were coherent in relation to the lineage tree, meaning that more closely related orthologs exhibited consistent changes in activity in relation to one another.

**Figure 4.**
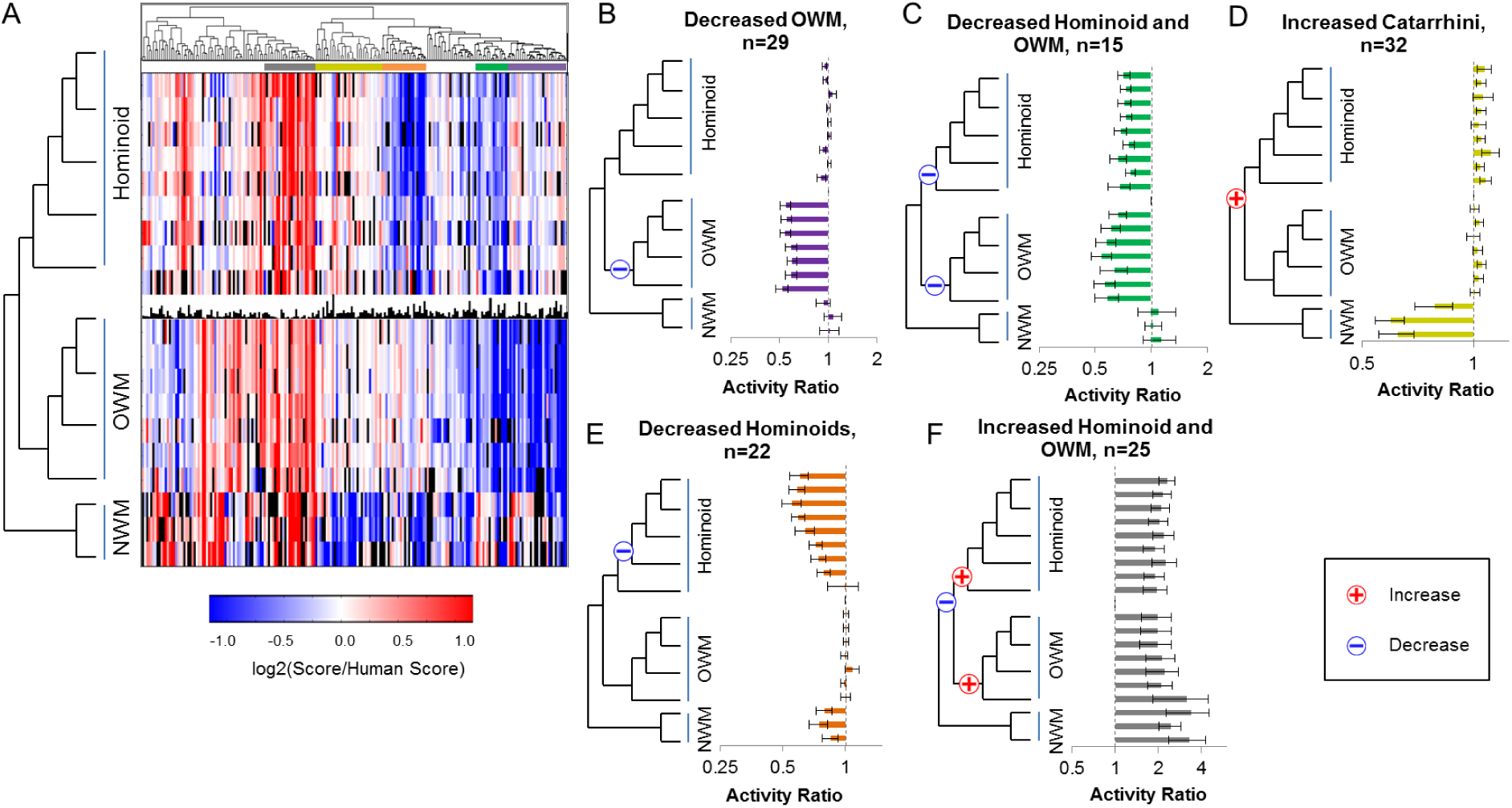
Common Patterns of Enhancer Modulation over the Primate Phylogeny. A) Functional scores for all enhancer tiles normalized to the MRCA to hominoids and Old World monkeys (N2). Black bar graph in the center contains the N2 score for each tile. Color bars above the heatmap indicate subsets of enhancer tiles exhibiting coherent patterns with respect to gain/loss of activity across the primate phylogeny, including: increased in human and hominoid (grey), increased in Catarrhini (yellow), decreased in hominoids (orange), decreased in hominoids and Old World monkeys (green), decreased in Old World monkeys (purple). B) The average score normalized to N2 for each species across the group of 29 enhancer tiles with decreased activity in Old World monkeys. Blue “-” indicates the timing of a loss of activity event. Error bars are one standard error. Dashed grey line at a ratio = 1. C) Same as Figure 4B for a group of 15 enhancer tiles with decreased activity in hominoids and Old World monkeys. D) Same as Figure 4B for a group of 32 enhancer tiles with increased activity in catarrhini. Red “+” indicates the timing of a gain of activity event. E) Same as Figure 4B for a group of 22 enhancer tiles with decreased activity in hominoids. F) Same as Figure 4B for a group of 25 enhancer tiles with decreased activity only in N2.

A first group (purple; **Figure 4B**) contains enhancer tiles that maintain activity in all orthologs except for Old World monkeys, which have consistently decreased activity relative to N2. This group contains 29 enhancer tiles (14%). A parsimonious explanation is that this group comprises tiles in which loss-of-activity events occurred on the branch between N2 and the MRCA to Old World monkeys.

A second group (green; **Figure 4C**) contains enhancer tiles with decreased activity in hominoids and Old World monkeys, relative to New World monkeys and N2. This group contains 15 enhancer tiles (7%). A potential explanation is that this group, the smallest of the five that we highlight in **Figure 4**, comprises tiles in which loss-of-activity events that occurred on both the branch between N2 and the MCRA of hominoids as well as on the branch between N2 and the MRCA of Old World monkeys.

A third group (yellow; **Figure 4D**) contains enhancer tiles with decreased activity restricted to the outgroup of New World monkeys. This group contains 32 enhancer tiles (16%), and is consistent with a single gain-of-activity event occurring between the MRCA to Simiformes (hominoids, Old World monkeys and New World monkeys) and N2, or alternatively a single loss-of-activity event on the branch leading to the New World monkeys.

A fourth group (orange; **Figure 4E**) contains 22 enhancer tiles (11%) with decreased activity in hominoids (lesser apes, great apes and humans) or only in hominids (great apes and humans) relative to N2. The most parsimonious explanation is a single loss-of-activity event along the branch from N2 to the MRCA of hominoids in some of these tiles and along the branch from the MRCA of hominoids to the MRCA of hominids in others (**Figure 4E**). This group is particularly interesting in that the human enhancer tiles, which are active based on ChIP-seq and our initial tiling experiment, have lower activity than some ancestral sequences as well as Old and New World monkeys. Looking more broadly, there are 35 enhancer tiles (17%) for which the human sequence exhibits significantly lower activity than the reconstructed N2 ortholog (p<0.05, 2-sample t-test). This bias towards reductions in activity relative to the ancestral N2 ortholog is not unique to human orthologs. Across the full dataset, 774 orthologs showed a significant reduction in activity compared to N2 ortholog while only 694 showed a significant increase (p=0.023, z-test two population proportions). This result suggests that the ancestral forms of regulatory sequences queried here tended to have greater activity than descendant sequences.

A fifth group (gray; **Figure 4F**) contains 25 enhancer tiles (12%) with decreased activity in N2 relative to all other orthologs. Given that this pattern would require 3 independent gains/losses for each enhancer, a simpler explanation may be that these represent low-quality reconstructions of the N2 sequence. As the most distant sequences from present-day orthologs, we expect N2 orthologs to be more likely to contain errors in reconstruction, which would in turn be expected to be biased towards reduction in activity. To test this, we extracted the marginal probability for each of our ancestral sequences from FastML. Overall, N10 (the MRCA to New World monkeys) had the lowest average marginal probability of 0.335, followed by N2 with an average marginal probability of 0.547 (**Supplemental Figure 4**). The remaining reconstructions all had marginal probabilities greater than 0.8. Next, we looked at the marginal probabilities for N2 from this group of 25 enhancers compared to the remaining 323 enhancers. This group had a significantly lower average marginal probability for the N2 reconstructions compared with the other enhancers (0.396 v 0.558, p=0.018, t-test for two independent means). This is consistent with the interpretation that the pattern observed for these 25 enhancers shown in **Figure 4F** is consequent to poor N2 reconstructions.

We examined whether tiles derived from the same enhancer peaks tend to fall within the same groups defined above. The 348 human-active enhancer tiles for which we tested additional orthologs derived from 233 candidate enhancers. Of these 233, 75 contained multiple tiles in our set, 8 of which had pairs of tiles that both fell within one of the five groups, which is significantly greater than expected by chance (p<1e-5, permutation test). Three of these eight pairs of enhancers were overlapping tiles, which can potentially narrow down the location of the causal mutation.

## Characterizing molecular mechanisms for enhancer modulation

We next explored the relationship between the sequence vs. functional evolution in enhancer activity across the primate phylogeny. As a starting point, we asked whether there was a correlation between the accumulation of sequence variation and the magnitude of change in functional activity for enhancer tiles. For every branch along the tree, we calculated the number of mutations between the mother and daughter nodes and the change in activity between the nodes. There was a significant, albeit modest, correlation between the number of mutations accumulated along a branch and the absolute change in functional activity (Spearman rho=0.207, p=4e-26) (**Figure 5A**). On average, each nucleotide substitution was associated with a 6.2% change in functional activity (slope of best fit line, with y intercept fixed to 1).

**Figure 5.**
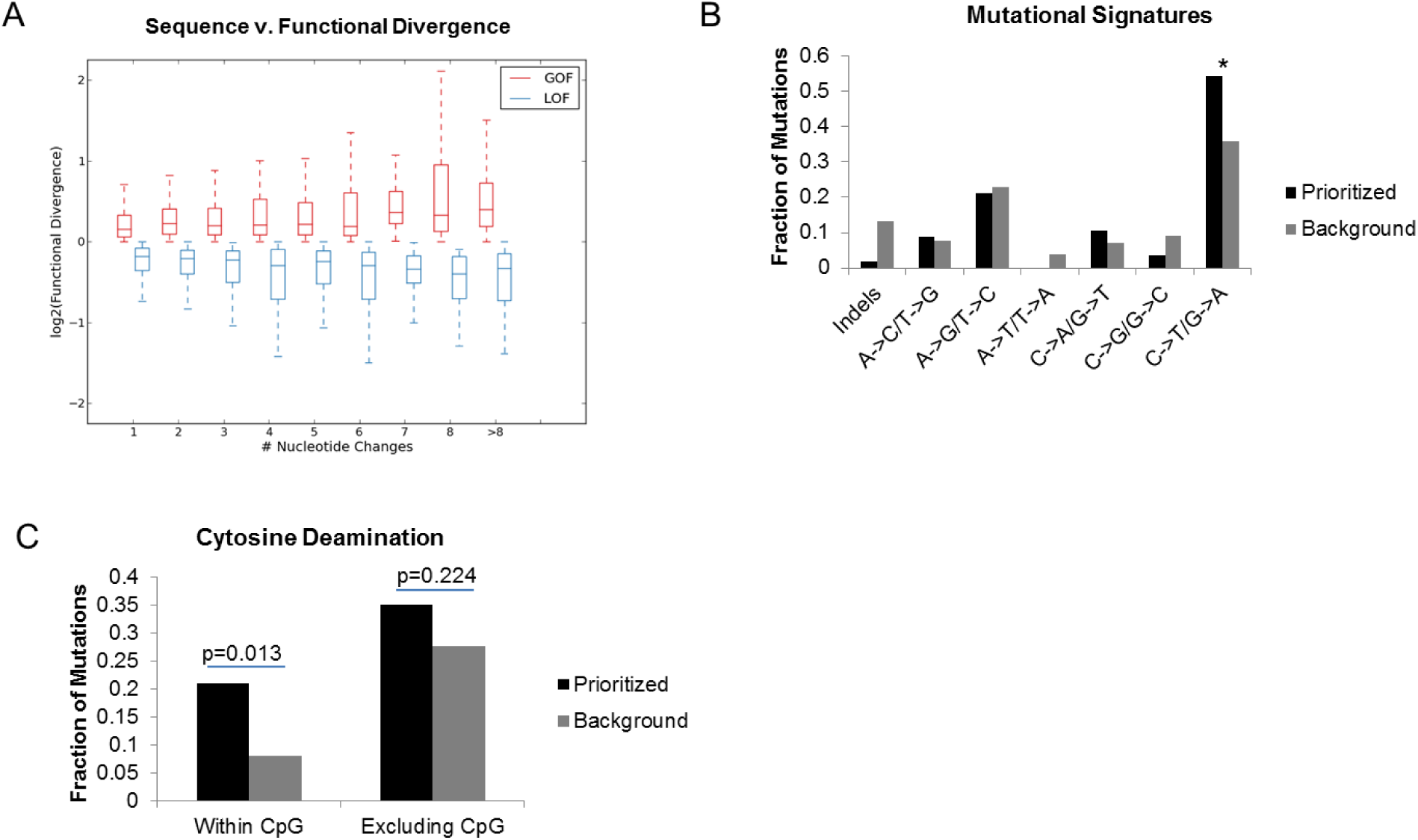
Molecular characterization of enhancer modulation. A) For every branch along the tree, we calculated the nucleotide and functional divergence. The number of nucleotide changes is on the x-axis and the log2 of the ratio of the daughter to ancestral functional score is on the y-axis. B) The fraction of indels, A→C, T→G mutations, A→G, T→C mutations, A→T, T→A mutations, C→A, G→T mutations, C→G, G→C mutations and C→T, G→A mutations in our set of 57 prioritized mutations (those associated with a significant functional difference) in black and 2,766 background mutations (those associated with a non-significant functional difference) in grey. Asterisk represents a Bonferroni-corrected p-value < 0.05 (Fisher’s exact test). C) The fraction of prioritized (black) and background (grey) mutations that are cytosine deamination events within CpGs and C→T, G→A mutations not disrupting a CpG.

We first asked whether different subsets of mutations were associated with functional changes. We focused on mutations that disrupt transcription factor (TF) motifs and asked whether these mutations were associated with changes in functional activity. For each enhancer tile, we identified all motifs associated with a TF expressed in HepG2 that were either lost or gained in at least one ortholog. We then ran linear regressions for the presence or absence of each TF motif against the functional scores for all orthologs, testing whether the mean slope of a TF from all enhancers was significantly different than zero using a two-sample t-test (**Supplemental Table 7**). The top two scoring TF motifs were E2F1, which was negatively correlated with activity *(i.e*. the presence of an intact motif is associated with decreased activity), and ATF2, which was positively correlated with activity *(i.e*. the presence of an intact motif is associated with increased activity). Both TFs play important roles in the liver [33,34], with E2F1 acting as a transcriptional repressor and ATF2 acting as a transcriptional activator. The E2F1 motif was disrupted in 60 enhancer tiles, and the ATF2 motif in 20 enhancer tiles. However, neither correlation was significant after a Bonferroni correction controlling for multiple hypothesis testing.

We next sought to prioritize specific mutations based solely on sequence vs. functional differences across the phylogeny. For each position along a given enhancer tile with a variant in at least one ortholog, we characterized each allele as ancestral (matching the MRCA of human and squirrel monkey, N2) or derived. We then performed Kolomogorov-Smirnov (K-S) tests at each position to test for association between allele status and functional scores, while applying a Bonferroni correction to account for the number of variants tested for each tile. Through this analysis, we identified a total of 57 mutations that we will refer to as “prioritized variants”, which correlate with the functional scores. We also generated a set of “background variants,” which did not correlate with functional scores (non-significant by the K-S test) (**Supplemental Table 8**). Within the 57 prioritized variants, there was a significant overabundance of C→T and G→A mutations over background (p=0.037, Fisher’s exact test, Bonferroni corrected) (**Figure 5B**). In order to test whether this effect is due in part to methylation, we looked at the subset of these C→T and G→A mutations, which disrupted a CpG. Cytosine deamination within a CpG accounted for 21% of our prioritized variants, compared to only 10% of background variants (p=0.013, Fisher’s exact test). When subtracting CpG deamination events, the enrichment for C→T and G→A mutations is no longer present (p=0.224, Fisher’s exact test), suggesting that CpG deamination events explain the observed enrichment.

### Discussion

While genome-wide studies demonstrate large-scale turnover of enhancers, the general molecular mechanisms underlying this turnover remain largely unexplored. In this study, we characterized modulation in the activity of hundreds of enhancer tiles throughout primate evolution, with nucleotide-level resolution. We first tried to characterize functional changes using computational tools, and although our tools were able to differentiate enhancers with low nucleotide identity (**Figure 2A**), they did not correlate well with our ChIP-seq-based predictions (**Figure 2C**) and performed poorly at predicting functional changes between evolutionarily similar sequences (**Figure 3B**). We therefore decided to test all sequences using STARR-seq, a reporter assay that experimentally measures regulatory activity for a library of sequences.

By testing all elements in the same *trans* environment (a single cell type), our experimental approach provided quantitative and directly comparable measurements, allowing us to measure functional differences between closely related sequences. However, this experimental approach assumes conserved trans-environments throughout the primate lineage. Previous studies have indeed noted that both the specificity of transcription factors for DNA and coactivators has remained highly conserved over much longer evolutionary time scales [35–38].

From both our computational predictions and functional scores, we note a low concordance with ChIP-seq based predictions (24%-37%). These numbers are similar to previous attempts to replicate biochemical predictions with high-throughput reporter assays [30,39,40], and there are plausible explanations for the difference. The ChIP-seq predictions were based on experimental data from primary liver samples from three individuals per species. Although most of the liver is composed of a single cell type, hepatocytes, there is still more diversity in such primary tissue than in the cell culture system we used for STARR-seq. Moreover, while we maintain a single *trans* environment for all of our orthologs, it is not an exact replica of primary liver. HepG2 cells are derived from a hepatocellular carcinoma, and likely have acquired changes during cancer development and immortalization. However, the fact that our enhancer tiles are both active in HepG2 cells (ChromHMM and STARR-seq) and in primary liver from humans (H3K27ac ChIP-seq) adds to our confidence that we are characterizing bona-fide enhancers.

Through hierarchical clustering of enhancer tiles normalized to human, we identified several functional groups. The largest group matched our ChIP-seq based predictions, with increased activity in humans and/or hominoids compared to other primate orthologs. We also identified a large group with decreased activity in Old World monkeys (concordant with three of the four ChIP-seq based predictions) and a third group with increase increased activity in Old World monkeys, or decreased activity in humans. The third group is the opposite of what we expected based on our ChIP-seq predictions. There are several possible explanations for this discordance in addition to the ones listed above. These regions may not have had ChIP-seq signal in rhesus, vervet, and marmoset if they were only active in a subset of liver cells (and therefore missed in a bulk-assay) or act as redundant (shadow) or poised-enhancer (and therefore lack active enhancer marks but are capable of activating transcription based on their sequence).

We next characterized evolutionary-functional trajectories for 206 of the enhancer tiles by normalizing all orthologs to the furthest ancestor, the MRCA between hominoids and Old World monkeys. We grouped these trajectories using hierarchical clustering, and identified several common patterns of modulation throughout the primate phylogeny. The most common patterns were tiles with a single loss of activity in Old World monkeys (n=29,**Figure 4B**), a single gain of activity in Catarrhini (or loss on the branch to New World monkeys) (n=32, **Figure 4D**), a single loss of activity in hominoids or hominids (n=22, **Figure 4E**), and two independent loss of activity events in hominoids and Old World monkeys (n=15, **Figure 4C**). We also identified a group of tiles with a decrease of activity unique to N2, without a clear parsimonious explanation (n=25, **Figure 4F**). Based on the lower marginal probabilities for the N2 reconstructions within this group, we propose that this subset of enhancer tiles may be due to incorrect reconstructions of the N2 sequence.

The group of enhancer tiles with decreased activity in hominoids may indicate sub-optimization or fine-tuning of enhancers. In total, 17% of our tiles showed a significant reduction of activity in human compared to N2, suggesting that reductions, without complete loss, of activity may in fact be a common phenomenon in primate enhancers. To determine whether sub-optimization was a general trend across the phylogeny, we calculated the number of enhancer tiles with significant increase or decrease of activity relative to N2. We identified significantly more tiles with decreases relative to N2 than increases. If we remove the cluster with reduced expression solely in N2, which we believe to be an artifact, the trend becomes more significant (768 vs 503, p<1e-5, z-test). All of these findings are concordant with high ancestral activity of present-day enhancers with subsequent loss to fine-tune activity along the phylogeny, at least for the enhancers that we chose to characterize here, which may be biased by the manner in which they were selected.

Ultimately, we wanted to look for general trends between sequence and functional divergence of enhancers throughout evolution. First, we looked at how the number of mutations accumulated along any branch on the tree correlates with the functional divergence along the branch. We found a modest, but significant correlation between sequence and functional divergence (Spearman rho=0.207, p=4.2e-26). While intuitive, to our knowledge, this is the first study quantifying the relationship between nucleotide changes occurring during evolution and experimentally-measured functional differences.

To further characterize mechanisms of mutations important in enhancer evolution, we utilized the high nucleotide identity between orthologs and reconstructed ancestral sequences to prioritize several variants, which were likely causal for functional divergence. While we first tried to prioritize variants based on transcription factor motif turnover, we did not find any significant motifs. Both ATF2 and E2F1 showed high correlations in our analysis, but neither was significant after multiple testing. Instead, we relied on prioritizing variants solely based on sequence content and functional scores, resulting in a list of 57 potentially causal variants. Of note, these 57 variants were enriched for cytosine deamination, particularly within CpGs, compared to variants that were not significantly associated with functional scores. Especially within closely related species, CpG deamination is a promising source of evolutionary novelty. Since spontaneous deamination of 5-methylcytosine (5mC) yields thymine and G-T mismatch repair is error prone, 5mC has a mutation rate four to fifteen-fold above background [41].

Besides its increased rate of mutation, there are multiple mechanisms by which CpG deamination may play a significant role in enhancer modulation. One mechanism is by introducing novel transcription factor binding sites or disrupting existing binding sites. In fact, Zemojtel *et al*. suggested that CpG deamination creates TF binding sites more efficiently than other types of mutational events [42]. CpG deamination may also alter enhancer activity by modifying methylation. Enhancer methylation has been correlated with gene expression, most frequently in cancer patients but also in healthy individuals [43]. Notably, enhancer methylation is both correlated with increased and decreased gene expression, possibly explaining why we see an enrichment of CpG deamination in both gain and loss of function events [44].

### Conclusion

In this study, we aimed to characterize general molecular mechanisms that underlie enhancer evolution. In order to do so, we conducted a large-scale screen of enhancer modulation with nucleotide-level resolution by combining genome-wide ChIP-seq with STARR-seq of many orthologs. We functionally characterized evolutionary-functional trajectories for hundreds of enhancer tiles, demonstrating a significant correlation between sequence and functional divergence along the phylogeny. We identify that many present-day enhancers actually have decreased activity relative to their ancestral sequences, supporting the notion of sub-optimization. We prioritized 57 variants, which correlated with functional scores, and found enrichment for cytosine deamination within CpGs among these potentially functional events. We propose that CpG deamination may have acted as an important force driving enhancer modulation during primate evolution.

### Supplemental Figures

**Supplemental Figure 1.**
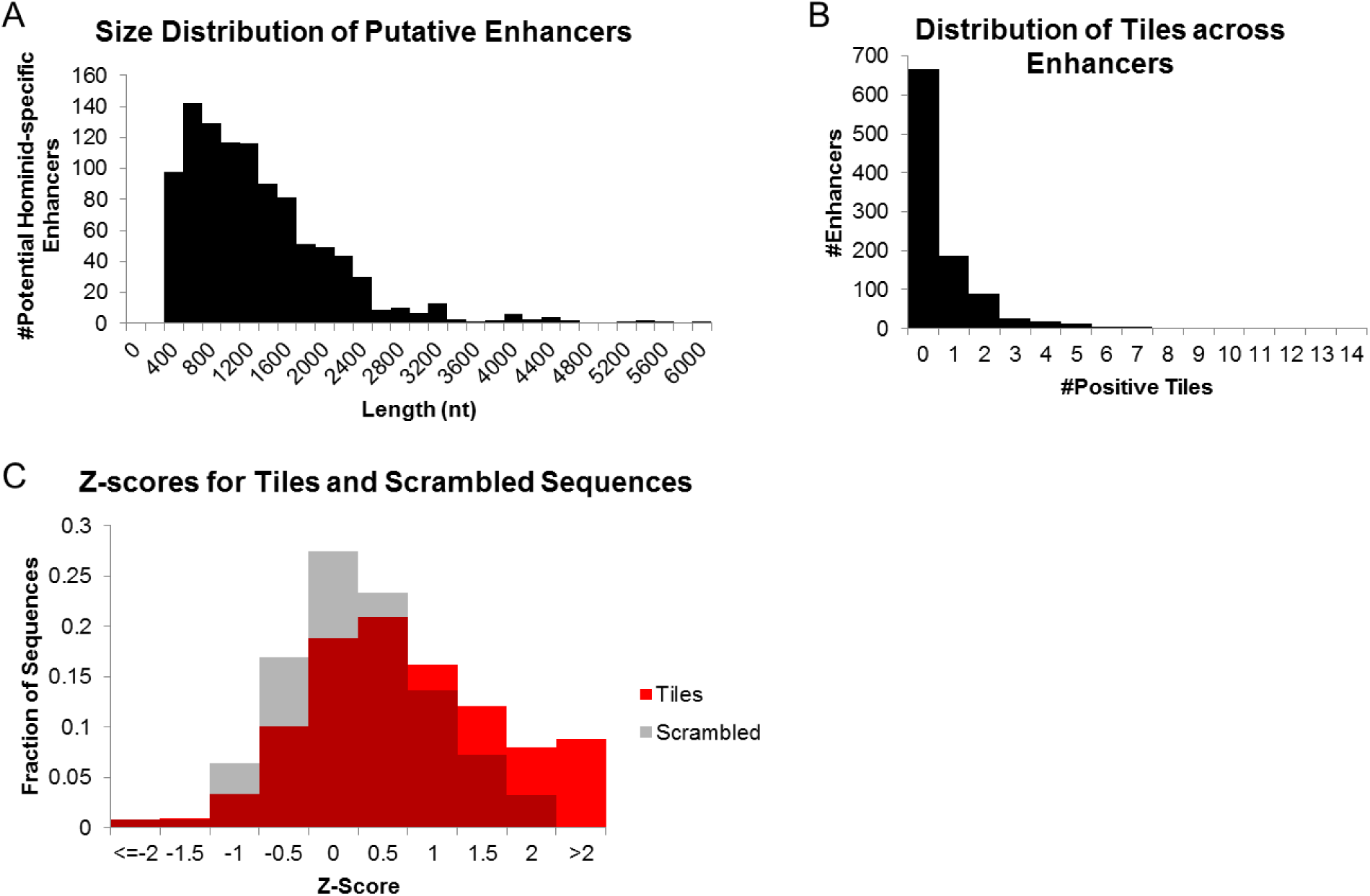
Tiling Across Large Enhancer Regions. A) Histogram of the size for each putative hominoid-specific gain-of-function enhancer defined by the intersection of H3K27ac ChIP-seq from primary tissue and HepG2 ChromHMM strong-enhancer calls. B) Histogram of the number of positive tiles (defined as a score two-standard deviations above the average negative control) per putative enhancer. C) Histogram of Z-scores for 6,724 tiles and 124 scrambled sequences.

**Supplemental Figure 2.**
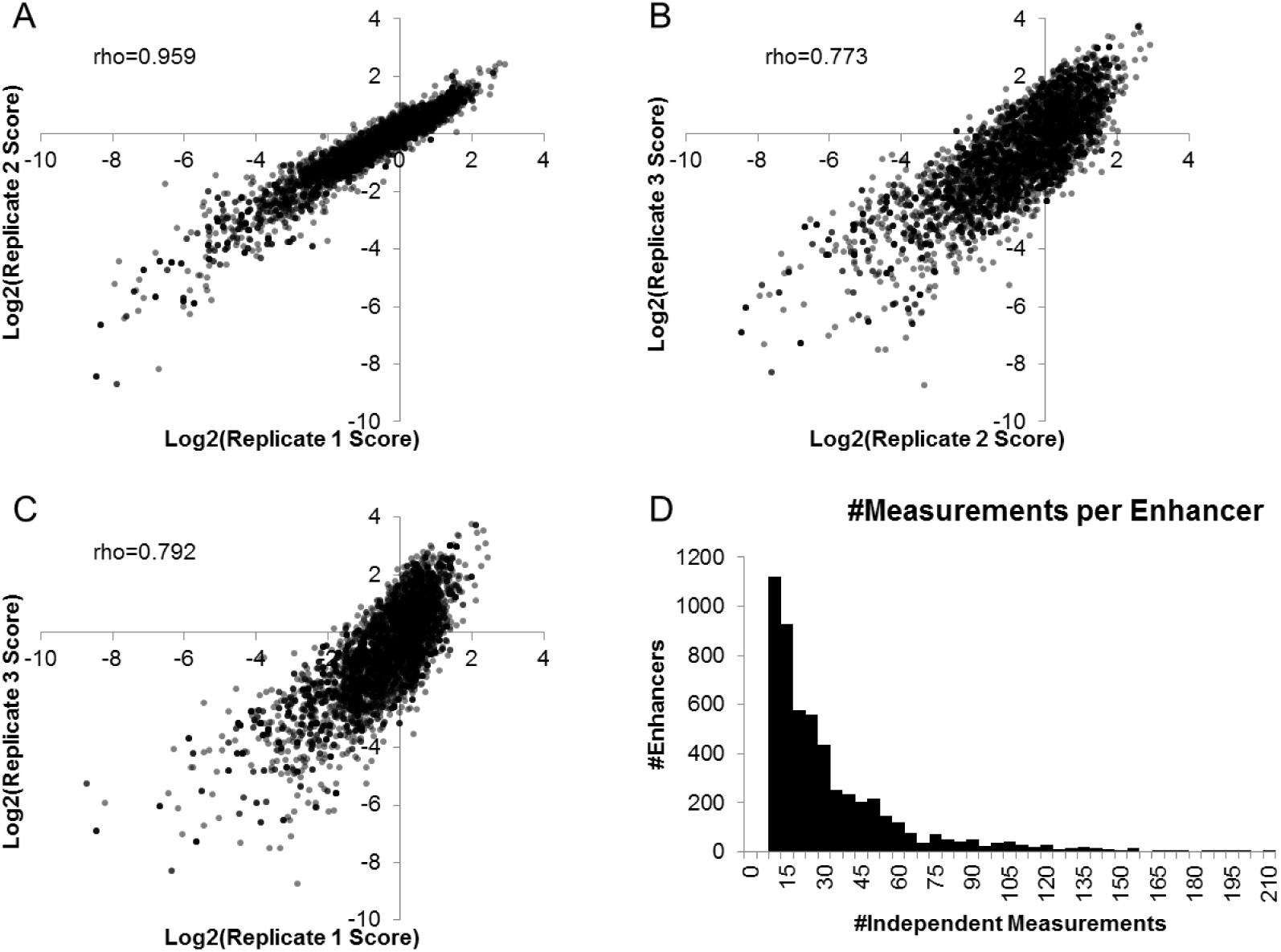
**Reproducibility of Functional Scores**. A) Spearman correlation between log_2_(normalized RNA/DNA) for biological replicates 1 and 2. B) Spearman correlation between replicates 2 and 3. C) Spearman correlation between replicates 1 and 3. D) Histogram of the number of independent measurements (barcodes) for each enhancer summed across all three replicates.

**Supplemental Figure 3.**
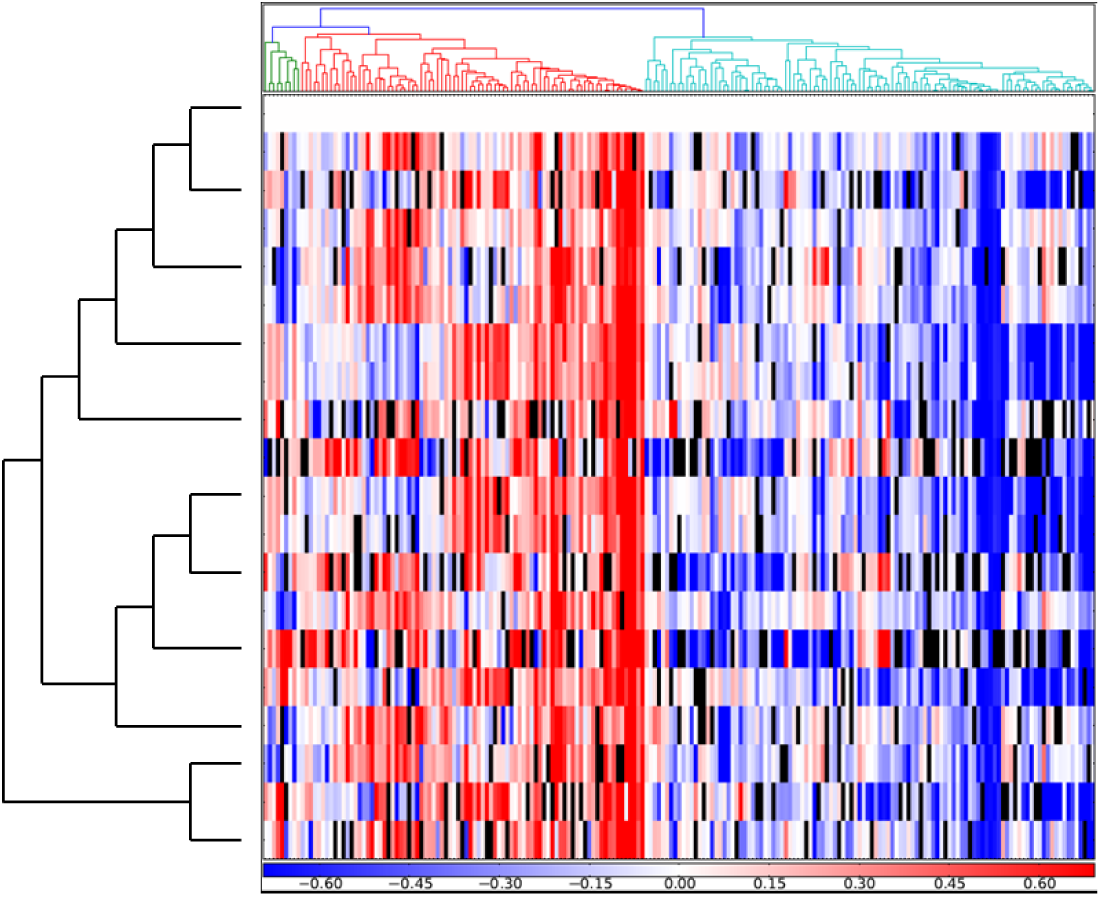
**Permuted Species’ IDs**. Species’ IDs were randomly permuted (seed=40) before normalization. Orthologs were hierarchically clustered and the same heatmap from Figures 2C and 3C was created.

**Supplemental Figure 4.**
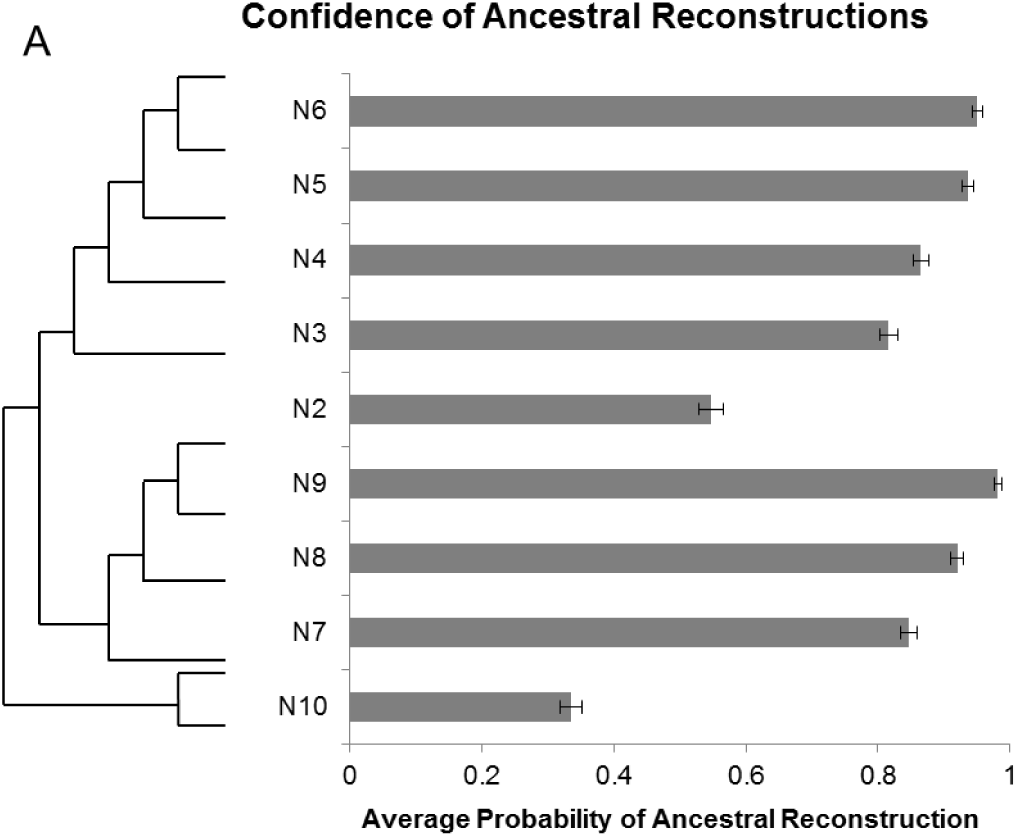
**Confidence of Ancestral Reconstructions**. The average marginal probability for each ancestral node across all 348 enhancers. Error bars indicate one standard error around the average probability for each node.

### Methods

#### Identification of potential hominoid-specific enhancers

We downloaded processed H3K27ac and H3K4me3 peak calls from Villar *et al*. [19]. Within each species, we called enhancers as H3K27ac peaks with a mean fold change ≥ 10 that were not within 1000bp of an H3K4me3 replicated peak. We converted all replicated H3K27ac peaks in rhesus, vervet and marmoset to hg19 coordinates using the UCSC liftover tool with a minimum match of 0.5. Villar *et al*. called the vervet peaks using the rhesus genome as a reference. We identified potential hominoid gain of function enhancers as predicted enhancers that did not have orthologous H3K27ac enrichment within 1kb from the summit in rhesus, vervet or marmoset. We converted the 10,611 gain of function enhancers back to the marmoset and rhesus genome with a minimum match of 0.9, with 6,862 having orthologs in the three genomes. We intersected our 6,862 GOF enhancers with ChromHMM strong enhancer calls in HepG2 using bedtools [45], resulting in a final set of 882 potential hominoid gain of function enhancers predicted to be active in HepG2.

#### Design and synthesis of tiles

For each potential hominoid gain of function enhancer, we defined end points by using the intersection of the H3K27ac peak and HepG2 ChromHMM strong enhancer call. For any intersections less than or equal to 200nt, we designed a 194bp tile around the center. For intersections with 200 ≤ length ≤ 400, we split the sequence into 3 overlapping fragments. For intersections > 400nt, we used 100bp sliding windows. We created negative controls from 800 tiles using uShuffle to create 200 dinucleotide shuffles each [46], and then picked the shuffled sequence with the fewest 7mers present in the original tile. We then synthesized all 10,544 tiles and 800 negative sequences as part of a 244K 230mer array from Agilent. The library was amplified from the Agilent array using the HSS_cloning-F (5’-TCTAGAGCATGCACCGG-3’) and HSS_cloning-R (5’-CCGGCCGAATTCGTCGA-3’) primers and cloned into the linearized human STARR-seq plasmid using NEBuilder HiFi DNA Assembly Cloning Kit [28]. The library was transformed into NEB 3020 cells and midi-prepped using the ZymoPURE Plasmid Midiprep Kit (Zymo Research).

#### Identification of active tiles

We transfected 5ug of our tiling library and 2.5ug of a puromycin expressing plasmid into three 60mm dishes, each with approximately 1.5 million HepG2 cells using Lipofectamine 3000 (ThermoFisher) according to manufacturer’s instructions. Twenty-four hours post-transfection, we selected cells with 1ng/mL puromycin for 24 hours. Forty-eight hours post transfection, we extracted DNA and RNA from the cells using the Qiagen AllPrep DNA/RNA Mini Kit (Qiagen). We treated RNA with the TURBO DNA-free Kit (ThermoFisher) and performed reverse transcription with SuperScript III Reverse Transcriptase (ThermoFisher). We amplified the cDNA using NEBNext High-Fidelity 2x PCR Master Mix with 5ul of RT reaction with primers HSS-F and HSS-R-pu1 in a 50ul reaction for three cycles with a 65°C annealing temperature. PCR reactions were cleaned with 1x Agencourt AMPure XP and eluted in 19ul (Beckman Coulter). We then performed a nested PCR using the whole purified cDNA reaction with primers HSS-NFpu1 (5’-CTAAATGGCTGTGAGAGAGCTCAGGGGCCAGCTGTTGGGGTGTCCAC-3’) and pu1R (5’-ACTTTATCAATCTCGCTCCAAACC-3’). DNA was amplified in one reaction using HSS-NFpu1 and HSS-R-pul (5’-ACTTTATCAATCTCGCTCCAAACCCTTATCATGTCTGCTCGAAGC-3’) with 1–2ug of DNA in a 50ul reaction and purified with 1.8x AMPure. We added barcodes and Illumina adaptors using Kapa HIFI HotStart Readymix in 50uL reactions with 1ul of previous PCR product with a 65°C annealing temperature and primers Pu1F-idx (5’-AATGATACGGCGACCACCGAGATCTACACACGTAGGCCTAAATGGCTGTGAGAGAGCTCAG-3’) and Pu1R-idx (5’-CAAGCAGAAGACGGCATACGAGATNNNNNNNNNGACCGTCGGCACTTTATCAATCTCGCTCCAAACC-3’) and sequenced on a 300 cycle NextSeq 500/550 Mid Output v2 kit with PE150bp reads. We aligned sequencing reads to the input library using BWA mem [47]. We then calculated RNA/DNA ratio for each sequence and defined active tiles as ones at least two standard deviations above the average negative sequence.

#### Design of orthologs and ancestral sequences

We identified all orthologs using the UCSC liftover tool with a minimum match of 0.9. For each sequence, we determined the longest ortholog, and set it to 194bp around the center. We then used LiftOver to identify the end points in other species. 348 of the 697 sequences were present through squirrel monkey, and we decided to use squirrel monkey as our outgroup moving forward. For ancestral reconstruction, we trimmed the hg38 phyloP 20way tree to the 11 species of interest and ran the Fastml heuristic [32]. We aligned each sequence with ClustalO to obtain a multiple sequence alignment [48], and then ran FastML (v3.1) with default settings on that alignment and the phyloP tree to create ancestral reconstructions.

#### Prediction of tiling and evolutionary results

We trained the gksvm-1.2 from Ghandi et al using the 500 top- and bottom-scoring lenti-MPRA sequences from Inoue et al as the positive and negative training sets, respectively, with default settings (Ghandi et al., 2014; Inoue et al., 2016). We used this model to predict scores for all tiles, and calculated the Spearman rho with our functional data. We next predicted scores for our positive human tiles and predicted negative orthologs from rhesus, vervet and marmoset and performed a two-sample t-test for each comparison. We calculated delta gkm-SVM scores by subtracting the predicted score of each ortholog from the predicted score of the human ortholog. We then predicted all eleven orthologs and nine ancestral nodes for all 348 enhancers.

#### Functional testing of orthologs and ancestral sequences

All orthologs and ancestral sequences were synthesized as part of an Agilent 230mer 244K array. We appended 5bp degenerate barcodes to each sequence by amplifying off the array with JK_R48_5N_HSSR (5’-CCGGCCGAATTCGTCGANNNNNCCATTGAGCACGACAGC-3’) and HSS_cloning-F (5’-TCTAGAGCATGCACCGG-3’). We then cloned the library into the STARR-seq vector in NEB 3020 cells, transfected into HepG2 cells, and prepared sequencing libraries as described above. Since some orthologs have very similar sequences, we aligned sequencing reads to our reference and only extracted error-free matches. We then calculated RNA/DNA ratios, and averaged across all barcodes for a given ortholog.

#### Molecular Characterization

We first looked to see whether the turnover of any transcription factor motifs correlated with functional scores. We ran FIMO to identify TF motifs from HOCOMOCO v9 that were lost or gained in at least one ortholog for each enhancer [49,50]. For one enhancer at a time, we ran a linear regression for the presence or absence of each TF motif against the functional scores of all orthologs tested. For each TF, we then tested whether the mean slope across all enhancers was equal to zero using a two-sample t-test and filtered for transcription factors with an FPKM ≥ 10, for a list of 134 motifs. We further filtered the list for TFs that turned over in at least 20 enhancers, further narrowing the list to 115 motifs.

We next looked to see whether any sequence mutations in an enhancer correlated with functional scores of the orthologs. For each enhancer, we performed a multiple sequence alignment using ClustalO. For each site along the enhancer (skipping the first to avoid alignment artifacts), we characterized the allele as ancestral or derived. For each site with a singleton derived allele in at least one ortholog, we performed a K-S test to see whether the allele associated with the functional scores. We then corrected the p-values for the number of sites along the enhancer that had derived alleles. For each site with only a single derived allele present in at least one species, we characterized the nucleotide change and summed the number of events over all enhancers. We calculated the Fisher’s exact p-value for each type of mutational event, using Bonferroni’s correction to adjust for multiple hypothesis testing. We then looked to see what fraction of C→T and G→A mutations disrupted CpGs, and calculated the Fisher’s exact p-values.

### Declarations

Ethics approval and consent to participate

Not applicable

### Consent for publication

Not applicable

### Availability of data and materials

All processed data from this study are included in this published article [and its supplementary information files’. Raw data are available from the corresponding author on reasonable request.

## Authors’ contributions

The project was conceived and designed by J.C.K. and J.S. J.C.K. and A.K. performed MPRA experiments. J.K. processed and analyzed data. T.D. wrote script for hierarchical clustering of enhancers. V.A. helped with tests for analyses. J.C.K. and J.S. wrote the manuscript. All authors read and approved the final version of the manuscript.

## Acknowledgement

The authors thank the Shendure lab (particularly J. Alexander and S. Kim), as well as S. Brothers for helpful discussions.

